# Early Reactivation of Emotion in ERPs to Neutral Retrieval Cues

**DOI:** 10.1101/594747

**Authors:** Holly J. Bowen, Eric C. Fields, Elizabeth A. Kensinger

## Abstract

Memory retrieval is thought to involve the reactivation of encoding processes. Previous fMRI work has indicated that reactivation processes are modulated by the residual effects of the prior emotional encoding context; different spatial patterns emerge during retrieval of memories previously associated with negative compared to positive or neutral context. Other research suggests that ERP indicators of memory retrieval processes, like the left parietal old/new effect, can also be modulated by emotion, but the spatial distribution and temporal dynamics of these effects is unclear. In the current study, we examined *when* emotion affects memory reactivation and whether that timing reflects processes that come before and may guide successful retrieval or post-recollection recovery of emotional episodic detail. While recording EEG, participants (N = 25) viewed neutral words paired with negative, positive or neutral pictures during encoding, followed by a recognition test for the words. Analyses focused on ERPs during the recognition test. In line with prior ERP studies, we found an early positive-going parietally distributed effect starting around 200 ms post retrieval-cue onset. This effect emerged for words that had been encoded in an emotional compared to neutral context (no valence differences), before the general old/new effect. This emotion-dependent effect occurred in an early time window, suggesting emotion-related reactivation is a precursor to successful recognition.

## Early Reactivation of Emotion in ERPs to Neutral Retrieval Cues

Memory for highly emotional positive and negative events, such as a surprise party thrown in your honor, or failing an important exam, is typically superior to memory for less emotional events, such as a conversation with your spouse about what to have for dinner (Bowen, Kark, & Kensinger, 2018; Brown & Kulik, 1977; Burke, Heuer, & Reisberg, 1992; Kensinger & Schacter, 2016). Empirical evidence also indicates that there are valence differences in memory. Young adults attend to and remember negative information better, and with more detail, than neutral or positive information (Baumeister, Bratslavsky, Finkenauer, & Vohs, 2001; Dewhurst & Parry, 2000; Kensinger, Garoff-Eaton, & Schacter, 2006; Kensinger & Schacter, 2006; Ochsner, 2000). For instance, young adults are better at identifying exactly which object (e.g., a yellow snake coiled up) they have seen in the face of similar distractors (e.g., a yellow snake stretched out) when the items are negative versus when they are positive (Kensinger, Garoff-Eaton, & Schacter, 2007). Understanding why memories of positive and negative events differ in this way is important for understanding the basic operation of human memory because so many of the experiences we remember have an affective tone.

Much of the research on these valence differences has focused on encoding (Kensinger, 2009). For example, research using functional magnetic resonance imaging (fMRI) during encoding of negative and positive subsequently remembered items indicates that arousing negative information engages more sensory processing regions (Mickley & Kensinger, 2008; Mickley Steinmetz & Kensinger, 2009) leading these negative items to later be remembered with more vivid detail (Kensinger, Addis, & Atapattu, 2011). Encoding of positive information, in contrast, engages areas in the prefrontal cortex that support conceptual processing and gist memory (Mickley & Kensinger, 2008).

In the current work, we focus on the effects of valence at retrieval. There is a long-standing hypothesis (James, 1980) that memory retrieval involves the reactivation or reinstatement of processes that were engaged when that information was encoded (Morris, Bransford, & Franks, 1977; Tulving & Thomson, 1973). Many modern theories of memory incorporate this idea, proposing that retrieval entails the recapitulation of distributed processes engaged during encoding (Alvarez & Squire, 1994; Bowen et al., 2018; Mather, Shafir, & Johnson, 2000; Moscovitch et al., 2005; Rolls, 2000), and empirical evidence has confirmed recapitulation of neural activity (Bowen & Kensinger, 2017b; Buckner & Wheeler, 2001; Hofstetter, Achaibou, & Vuilleumier, 2012; Kark & Kensinger, 2015, 2019; Rugg, Johnson, Park, & Uncapher, 2008; Wheeler, Petersen, & Buckner, 2000). For example, fMRI studies have demonstrated sensory-specific reactivation such that stimuli previously encoded visually or auditorily activate the visual and auditory cortex, respectively, at retrieval (Gottfried, Smith, Rugg, & Dolan, 2004; Nyberg, Habib, McIntosh, & Tulving, 2000; Slotnick, 2004; Wheeler & Buckner, 2004; Wheeler et al., 2000). Content-specific reactivation has also been demonstrated in the fusiform gyrus during the retrieval of information previously encoded with a face (Hofstetter et al., 2012; Skinner, Grady, & Fernandes, 2010).

This type of evidence supports the theory that neural processes at retrieval are similar to those engaged when the information was originally encoded, and these effects reflect reactivation of the associated information. To this end, our prior fMRI work has shown that neutral retrieval cues can reactivate some of the emotion-related properties associated with the prior encoding experience. Successful memory for neutral words (Bowen & Kensinger, 2017a, 2017b) and line drawings (Kark & Kensinger, 2015, 2019) is associated with greater recapitulation in regions including those within the ventral visual stream to a greater extent for items previously encoded in a negative compared to a neutral or positive context. This is true even though the emotional content is no longer present at the time of retrieval (for a review see Bowen, Kark, & Kensinger, 2018).

### ERP evidence for neural recapitulation

The reactivation evidence presented above was from fMRI studies, but there is also a literature assessing content-dependent electrophysiological correlates and the time course of memory retrieval using event related potentials (ERPs). Johnson and colleagues (Johnson, Minton, & Rugg, 2008) examined recollection of words that differed according to their prior encoding history by either incorporating the word in a sentence or superimposing it on an image. They found that ERPs elicited by words previously encoded in a sentence compared to an image were more positive-going, and the effect was enhanced for recollective memory compared to familiarity responses. The discrepancy between the two waveforms began to emerge in anterior sites approximately 300 ms post stimulus continuing until about 1000 ms. These content-dependent ERPs had an onset as early as the general left parietal old/new effects demonstrating that content-dependent neural activity during retrieval can occur in a timeframe which suggests it is guiding the retrieval process, and specifically recollection, of episodic information. A second effect beginning around 800 ms post stimulus onset distributed bilaterally over posterior sites and opposite in polarity to the early effect might reflect monitoring or maintenance of information retrieved.

In a more recent study, Johnson and colleagues (Johnson, Price, & Leiker, 2015) employed multivariate pattern analysis to examine the timing of reactivation of encoding processes at retrieval. They found that reactivation was co-occurring with ERP left parietal old/new effect (~500 ms post stimulus onset). The reactivation effects were also positively correlated with behavioral accuracy suggesting that reinstatement was capturing and is involved in successful retrieval. Effects that occurred later in the time course were consistent with the maintenance or monitoring and post-retrieval processes. Finally, Waldhauser and colleagues (Waldhauser, Braun, & Hanslmayr, 2016) found evidence that early sensory reactivation is actually necessary for episodic memory. Applying rhythmic transcranial magnetic stimulation to lateral occipital cortex during retrieval cues disrupted desynchronization in the α/β band thereby decreasing spatial source memory for those cues.

The novel findings from these studies is the temporal characterization of reactivation, such that reactivation is related to and potentially guides episodic retrieval success early in the time course, but that reactivation also contributes to post-retrieval processing and stimulus evaluation.

### Effects of emotion at encoding on ERPs at retrieval

A few ERP studies have examined the effects of emotion on memory retrieval. These studies all use a similar design: Items (words or pictures) are initially encoded in emotional contexts. At retrieval these items are presented without the previously presented context. The key question is whether the ERPs elicited at retrieval differ based on the emotional properties of the encoding context.

Maratos and Rugg (2001) presented sentences that contained neutral critical words in a broader context that was either negative or neutral. When the neutral critical word was re-presented at retrieval, there was a left parietal old/new effect beginning by 500 ms for all old items when compared to new items, but this effect was larger for words encoded in a negative context. There was also a later (after 800 ms) right frontal positivity that was larger to words that had been encoded in a negative context. Interestingly, these valence effects disappeared on both components in a second study when at retrieval participants were required to make a source judgment about the emotionality of the prior encoding context. The authors suggested that participants may have been more likely to spontaneously retrieve the source context for items encoded in negative contexts, and that this effect therefore disappears when the task requires retrieval of source context for all items.

Smith, Dolan, and Rugg (2004) report a similar study using photos: neutral objects were presented on neutral or emotional backgrounds. In contrast to the Maratos and Rugg study, the valence of the context primarily showed effects at frontal sites after 800 ms, with only weak evidence for earlier effects. These effects held whether source judgments were required or not. Jaeger, Johnson, Corona, and Rugg (2009) used a very similar picture-based paradigm to examine the effects of study-test delay (see also Jaeger & Rugg, 2012). However, in the short-delay condition—a near replication of Smith et al. — more robust differences between items encoded in negative versus neutral contexts emerged early, around 200 ms at frontal sites, and continued at left central sites from around 300 ms through the end of the recorded epoch (1100 ms).

It is difficult to draw firm conclusions from this literature. On the one hand, it clearly suggests that traditional indicators of memory retrieval processes like the left parietal old/new effect can be modulated by emotion. On the other hand, the spatial distribution and reports of how early these emotion effects emerge in the retrieval process are variable despite similar paradigms across studies.

One issue with evaluating the differences across studies is the statistical approach employed. These studies analyzed data using factorial ANOVAs with three spatial factors to explore effects across different regions of the scalp. Although this approach has long been common in ERP research, the field has recently become more aware that the inclusion of such spatial factors leads to significant multiple comparisons issues (see discussion in Fields & Kuperberg, 2018; Luck & Gaspelin, 2017). Although the later effects (after 800 ms) appear robust in these studies, early effects often emerged only in complex interactions with spatial factors and were not significant at levels that survive multiple comparisons correction.

In addition, most studies have only compared negative contexts to neutral contexts, which leaves open important questions about the effects of valence. Only Smith et al. (2004) included both positive and negative contexts during the encoding phase. That study showed effects of encoding in emotional contexts versus neutral, but showed no differences between positive and negative contexts. This is in contrast to the fMRI literature reviewed above, which has shown greater neural recapitulation for stimuli encoded in negative contexts. It is unclear whether these differences are due to differences in the stimuli or paradigms employed, or whether they reflect differences in the neural processes that fMRI and ERP are sensitive to.

### The present study

In the current work, we employed a paradigm known to elicit greater recapitulation for negative versus positive contexts in fMRI (Bowen & Kensinger, 2017a, 2017b). Neutral words were paired with an unrelated neutral, positive, or negative picture during memory encoding. Each participant had a customized set of stimuli based on their individual valence and arousal ratings from a larger stimulus set, which ensured that participants were indeed seeing stimuli that they deemed negative, positive and neutral. At retrieval, ERPs were recorded to presentation of the neutral words presented in isolation. We analyzed data using a mass univariate approach (similar to the way fMRI data is commonly analyzed); recent work has highlighted the value of this approach for controlling multiple comparisons issues while maintaining flexibility to find effects when the precise spatial and temporal distribution is not known in advance (Fields & Kuperberg, 2018; Groppe, Urbach, & Kutas, 2011). Given findings in the literature reviewed above, we employed analysis parameters that allowed us to identify effects at any scalp location and in both early and late time windows.

Our goal was to answer two key questions. First, we asked whether, consistent with the fMRI literature, negative and positive valence would show differing effects at retrieval, or whether, like the Smith et al. studies describe above, emotional contexts would modulate retrieval when compared to neutral contexts without differences between positive and negative. Second, we examined the timing of these effects. Early modulation by previous emotional context would suggest that emotion at encoding affects processes that guide successful retrieval (cf. Johnson et al., 2008; Waldhauser et al., 2016). If such effects are only present in later time windows, this would suggest that emotion effects may emerge only after successful retrieval.

## Methods

### Participants

All interested participants completed an extensive medical screening questionnaire to assess past and current medical conditions such as psychological disorders, head trauma or injuries, and medications. Reasons for ineligibility included currently taking antidepressant or anxiolytics, head trauma (e.g., concussion) in the last 2 years, seizures, diabetes, learning disorder (e.g., dyslexia, ADHD). We also required that participants be right handed and native English speakers. All procedures were approved by the Boston College Institutional Review Board and all participants gave informed consent. A total of 41 participants were tested, the majority of which were from the Boston College community with some additional participants recruited through a posting on craigslist.org. Participants were either paid $15/hour or given partial course credit. One participant withdrew prior to completion due to a headache, five participants were excluded because of technical errors or equipment failure, four were excluded because of poor behavioral performance (e.g., overall performance lower than chance or too few responses for reliable ERP analysis), and six datasets were unusable due to excessive artifact in the EEG (greater than 25% of trials lost to artifact—see artifact correction and rejection procedures below). This left 25 eligible participants (N = 8, male) with an average age of 22 years (range 18-32 years). All eligible participants completed the Beck Depression Inventory (Beck, Epstein, Brown, & Steer, 1988) and scored below 11 (*M* = 1.96, SD = 2.39).

### Stimuli

Three different types of stimuli were used in the study: faces, scenes and words. The stimuli were gathered from a number of sources as detailed below. During encoding, stimuli stayed on screen for 3.5 seconds and during retrieval until a response was made. Each stimulus was followed by a fixation that was jittered 500-4000 ms in both phases of the experiment.

#### Faces

Forty-two face stimuli were selected from the NimStim (http://www.macbrain.org/resources.htm) Set of Facial Expressions (Tottenham et al., 2009). These faces range in emotional expression, valence and arousal, gender, race and ethnicity.

#### Scenes

Fifty-seven total images of scenes which contained no perceptible faces were used. To meet this requirement some images were taken from the International Affective Picture System (IAPS; (Lang, Bradley, & Cuthburt, 2008) and some from various sources on the internet. The scenes depicted in the images ranged in content as well as emotional valence and arousal.

#### Words

Words were selected from the published dataset (Warriner, Kuperman, & Brysbaert, 2013) consisting of 13,916 words that include valence and arousal ratings. The large list was reduced to six-letter neutral words, and then further restricted to 384 words using a random number generator (M_valence_ = 5.71, SD_valence_ =1.64, range = 5.0 – 6.9; M_arousal_ = 3.41, SD_arousal_ = 2.16, range = 2.27-3.96). Words were then split into 8 groups of 48 words. The 8 lists did not differ on average valence or arousal ratings, *F*(7, 383) = .85, *p* =. 55 and *F*(7, 383) = .65, *p* = .72, respectively.

### Procedure

Before being scheduled to come to the lab on Boston College campus, all interested participants completed the medical screening questionnaire. Once eligibility was determined, participants were asked to complete an online questionnaire via surveymonkey.com to rate the valence and arousal of all the scene and face stimuli and were then scheduled to come into the lab for a single two-hour session. The online questionnaire had participants rate all 42 faces and 57 scenes on valence and arousal. The purpose was to create an individualized subset of these images (4 scenes and 4 faces of each valence; 24 images total) as experimental stimuli; the specific images selected for each participant were the 8 negative and 8 positive images that were given the highest arousal ratings during the initial ratings task, along with 8 neutral rated images. This ensured that participants were viewing images that indeed were emotional in their opinion. Valence and arousal from one participant are not included in analyses below due to experimenter error.

When participants came to the lab they provided informed consent, demographic information, and completed the Beck Depression Inventory (BDI). They were explained the EEG procedure, viewed the EEG equipment, and were encouraged to ask questions before proceeding. After the EEG set-up was completed, participants were given instructions about the task and completed a practice of six encoding and twelve retrieval trials.

Participants completed four alternating encoding-retrieval blocks with a short, filled retention interval separating the two phases. During the encoding phase (48 trials per block), participants viewed a word and an image on the screen together. They were instructed to try and remember the word for the upcoming memory test and indicate via button press whether the image was of a face or a scene. The encoding block was followed by ten multiple choice math problems (e.g., 2+5, 6 × 2). During the retrieval phase, participants viewed only the words and were asked to make a ‘remember’, ‘know’, ‘guessing’ or ‘new’ judgment. There were 48 target items and 48 lures (96 items total) during each retrieval block. Participants were given the remember and know instructions (Rajaram, 1993), were encouraged to respond “guessing” if they had no inclination as to whether the word was old or new, and to respond “new” if they had not seen the word during the encoding stage. ERPs were recorded during both the encoding and retrieval phases of the study. Participants were given a short break after the second block of trials. After the study was complete, participants were compensated, debriefed, and thanked for their participation.

### ERP Acquisition and Processing

Data was collected using a BioSemi ActiveTwo EEG system and ActiView v6.05 EEG acquisition software (http://www.biosemi.com/). The EEG was recorded from 32 Ag/AgCl electrodes in an elastic cap placed according to the international 10-20 system^1^. In addition, electrodes below and to the left of the left eye and above and to the right of the right eye were recorded to monitor for blinks and eye movements, and electrodes on each mastoid were recorded to serve as the reference. The EEG signal was amplified, filtered online with a low pass 5th order sinc response filter with a half-amplitude cutoff at 104 Hz, and continuously sampled at 512 Hz.

### Data Processing and Analysis

EEG and ERP data processing were conducted in EEGLAB v14.1.1 (https://sccn.ucsd.edu/eeglab/index.php; Delorme & Makeig, 2004) and ERPLAB v6.1.4 (https://erpinfo.org/erplab; Lopez-Calderon & Luck, 2014).

The EEG was first referenced to the average of the two mastoid electrodes. For each segment of continuous EEG, we subtracted the average voltage of the entire segment, then applied a high-pass 2nd-order Butterworth infinite impulse response filter with a half-amplitude cut-off of 0.1 Hz (Kappenman & Luck, 2010; Tanner, Morgan-Short, & Luck, 2015). We then extracted segments from 200 ms before to 1100 ms after all events of interest.

For the purposes of independent components analysis (ICA), the mean voltage was removed from each segment at each electrode (Groppe, Makeig, & Kutas, 2009). Segments containing significant artifact that was not neural, ocular, or muscular in origin were identified via visual inspection and excluded from the data submitted to ICA. Independent components were calculated from the remaining segments using the extended infomax algorithm (Lee, Girolami, & Sejnowski, 1999) as implemented in EEGLAB. Independent components corresponding to ocular activity (blinks and saccades) were identified by visual inspection and removed. This consisted of either 2 or 3 independent components for each subject.

After independent components were removed, all segments were re-baselined to the mean of the pre-stimulus period. Segments with remaining artifact were identified via artifact detection algorithms implemented in ERPLAB. The parameters of these algorithms (e.g., voltage thresholds) were tailored to each subject via visual inspection of the data, but were consistent across all conditions within each subject. Trials containing a blink or large saccade within the first 200 ms of a trial were rejected even if the artifact was corrected via ICA, as these trials may have a delayed neural response due to the eyes being closed or averted during stimulus presentation. Rejection rates ranged from 2% to 24% across subjects with an average of 11% and did not differ significantly across valence conditions.

After ICA based correction and artifact rejection, remaining trials were averaged within conditions of interest to form ERPs.

### ERP analysis

Statistical analysis of ERP was conducted via the Mass Univariate Toolbox (Groppe et al., 2011) and Factorial Mass Univariate ERP Toolbox (https://github.com/ericcfields/FMUT/wiki). We used a cluster-corrected mass univariate approach (Groppe et al., 2011; Maris & Oostenveld, 2007). Briefly, this approach consists of conducting an ANOVA independently at each time point and electrode of interest. Clusters are identified as adjacent time points/electrodes with effects surpassing a threshold, and all *F*-values in the cluster are summed to form a cluster mass statistic. A permutation approach is used to estimate the null distribution for this statistic, which is then used to calculate a *p*-value for each cluster. Recent simulation work has shown that this approach can provide power that compares well to traditional mean amplitude based approaches while providing significantly greater flexibility (Fields & Kuperberg, 2018).

Because the relevant literature suggested effects may occur at frontal and posterior sites and in both early and late time windows, the cluster mass test was conducted at all time points from 200 to 1000 ms at all 32 scalp electrodes. The *F*-value corresponding to *p* = .05 in an uncorrected parametric test was used as the threshold for cluster inclusion and electrodes within approximately 7.5 cm of each other (assuming a head circumference of 56 cm) were considered neighbors. 100,000 permutations were performed for each test. Statistical analysis was conducted on data that was first low-pass filtered at 10 Hz (half amplitude cut-off, 2nd-order Butterworth IIR filter) and downsampled to 128 Hz (Groppe et al., 2011; Luck, 2014). Cluster significance was evaluated at α = 0.05.

## Results

### Behavioral Data

#### Stimulus arousal ratings

Arousal ratings for the experimental stimuli were submitted to a 3-way (valence: negative, positive, neutral) repeated measures ANOVA. There was a significant effect of valence, *F*(2, 46) = 113.08, *p* < .001, ω_p_^2^ = .82. This interaction was followed-up with paired samples *t-* tests which indicated that negative stimuli (*M* = 6.43, *SD* = .66) were rated as more arousing than positive (*M* = 5.46, *SD* = .86), *t*(23) = 6.06, *p* < .001, *r* = .78 and positive were more arousing than neutral stimuli (*M* = 3.94, *SD* = .27), *t*(23) = 8.24, *p* = .001, *r* = .86.

#### Accuracy and reaction time

Average hit rates^2^ were calculated for stimuli previously encoded with negative, positive and neutral images and submitted to a 3-way repeated measures ANOVA. The main effect of emotion was not significant, *F*(2, 48) = 2.41, *p* = .10, ω_p_^2^ = .05, indicating the hit rate for items previously encoded with negative images, *M* = .81 (*SE* = .02), positive images, *M* = .83 (*SE* = .03), and neutral images, *M* = .79 (*SE* = .03) did not statistically differ. Since new stimuli presented at retrieval were never associated with a particular valence there is only a single false-alarm rate. Overall corrected recognition (hit rate minus false alarm rate) was *M* = .31 (*SE* = .04). Median reaction times for recognition of stimuli previously encoded with negative, positive and neutral images were calculated and submitted to a 3-way repeated measures ANOVA. Again, the main effect of emotion was not significant, *F*(2, 48) = .25, *p* = .78, ω_p_^2^ = -.03, indicating memory-related reaction times to items encoded with negative, *Md* = 1321 (*SE* = 77), positive, *Md* = 1303 (*SE* = 64), and neutral images *Md* = 1317 (*SE* = 60) did not statistically differ. The median reaction time for false alarms was *Md* = 1554 (*SE* = 102).

#### Proportion of responses

Table 1 displays the proportion of remember, know, guess and new responses for target stimuli within each level of valence and new items. To align with the ERP analyses described below, we were most interested in whether the proportion of items correctly recognized as old (i.e., given either a remember or know response) would vary by emotion. A 3-way repeated measures ANOVA indicated that this proportion did not vary for negative (*M* = .64, *SD* = .16), positive (*M* = .64, *SD* = .17), or neutral encoding contexts (*M* = .66, *SD* = .18), *F*(2, 48) = .704, *p* = .50, ω_p_^2^ = -.01.

**Table 1.** Proportion of remember, know, guess and new responses to target and distractor stimuli given an initial “old” response within each valence category.

| Response | Negative Target | Positive Target | Neutral Target | Distractor |
| --- | --- | --- | --- | --- |
| Remember | .45 (.03) | .44 (.03) | .41 (.04) | .07 (.02) |
| Know | .19 (.02) | .21 (.03) | .24 (.03) | .14 (.02) |
| Guess | .18 (.02) | .17 (.03) | .16 (.02) | .26 (.03) |
| New | .15 (.02) | .14 (.02) | .16 (.02) | .50 (.04) |

Note. Distractor (i.e., new) items did not vary by emotional context. Standard error in parentheses.

### ERP Data

Visual inspection of the EEG and ERPs at encoding suggested that subjects were variable from trial to trial in terms of when they were looking at the word versus the picture (which were presented simultaneously). Since neural processes must be aligned in time across trials to form meaningful ERPs, and because the key research questions in the present study were about neural processes at retrieval, the data recorded at encoding is not considered further.

ERPs were collapsed across Remember/Know (considered together as an “Old” response) and Guess/New (considered together as a “New” response) to ensure enough trials (> 15 for each subject) went into each average to provide stable ERPs and reasonable power. Visual inspection revealed that effects of valence were similar for Remember and Know trials, but the Know trials were noisier due to smaller bin sizes (see Table 1). Separate results for each of the four judgments can be seen in the supplementary materials (available on the Open Science Framework https://osf.io/jyd5c/).

Figure 1 shows the effects of valence at retrieval. For trials given a Remember or Know judgment, words that had been paired with a positive or negative picture began to show a more positive amplitude than those previously paired with a neutral picture by 200 ms after word onset. This emotion effect had a parietal distribution and continued to the end of the epoch. As shown in Figure 2, a cluster beginning at 213 ms was significant in the statistical analysis (*p* = .009). To assess our first research question, we followed-up this main effect by calculating all pairwise comparisons using the same analysis parameters. These contrasts showed that words encoded with a positive or negative picture both elicited a larger positivity than those encoded with neutral pictures (*p* = .011 and *p* = .006, respectively). The response to positive and negative was not significantly different (*p*s > .50).

**Figure 1.**
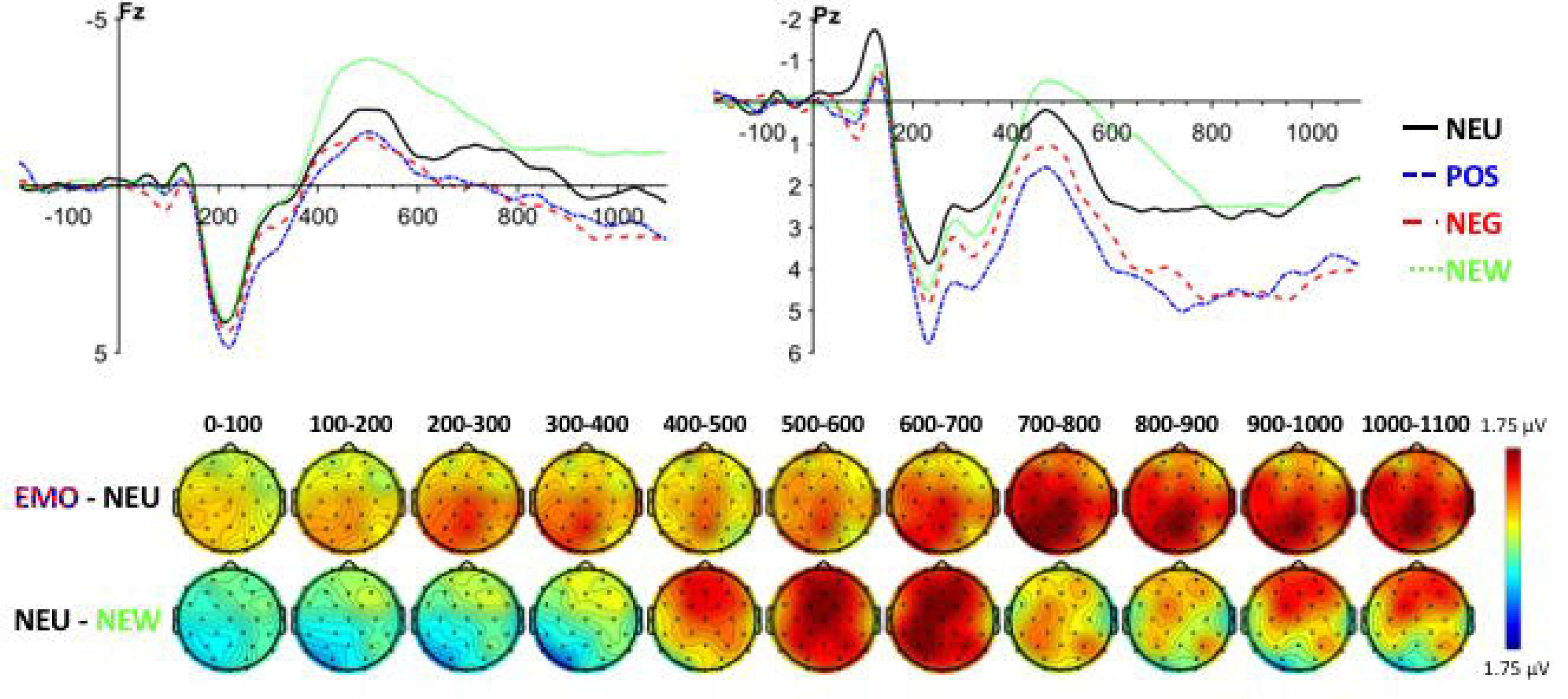
ERP waveforms show the response at retrieval to words previously encoded with neutral, positive, and negative photos and given a remember or know judgment, as well as new words given a guess or new judgment. Scalp maps show the emotion (collapsed across positive and negative) minus neutral effect and the old/new (neutral – new) effect. ERPs are filtered at with a 10 Hz half-amplitude low pass filter, matchiing the data used for statistical analysis (see Methods).

**Figure 2.**
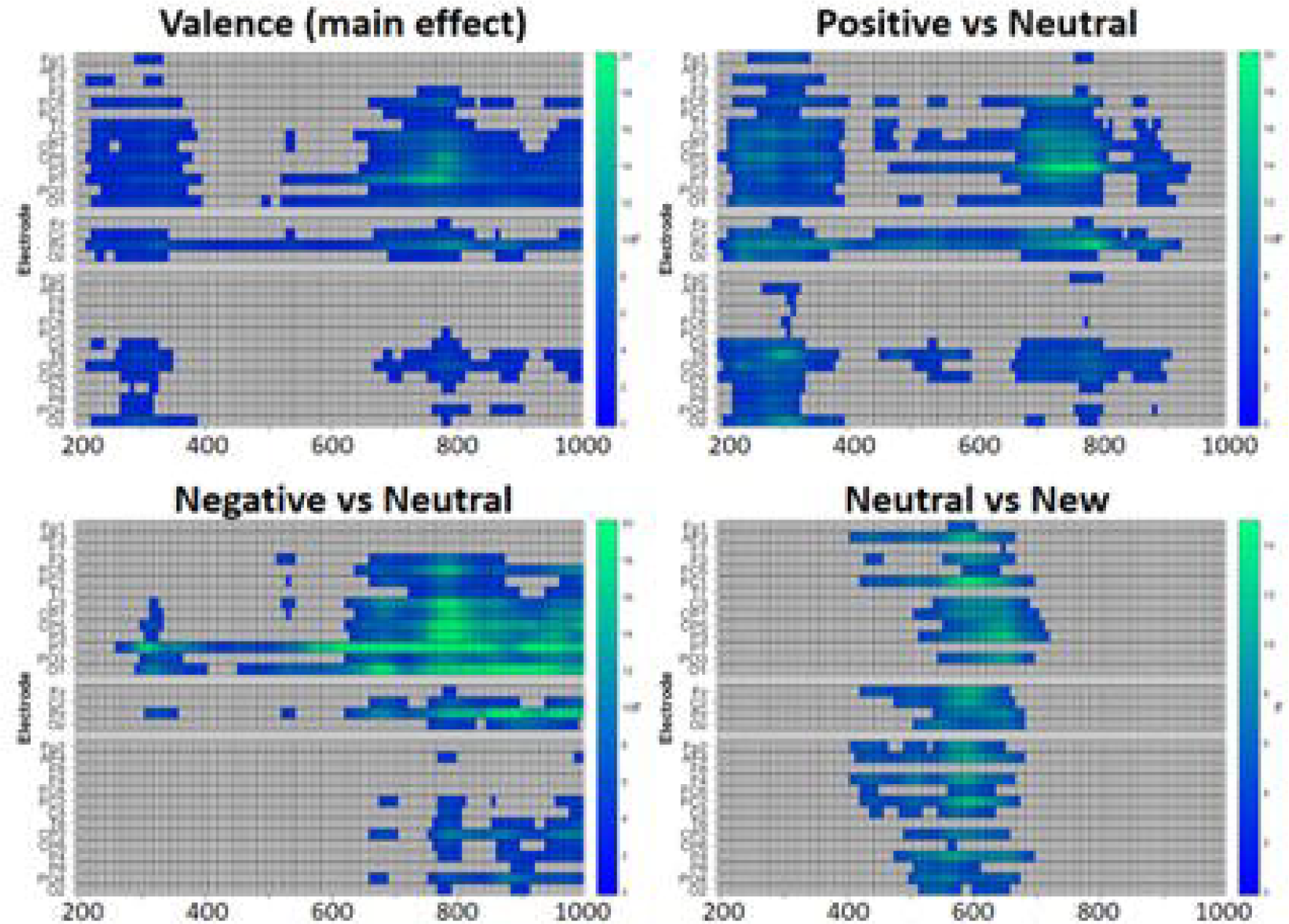
Results of the cluster-corrected statistical analysis. Positive, Negative, and Neutral refer to the ERP at retrieval to words encoded with a picture of each valence and given a remember or know judgment. New refers to words presented at retrieval that were not previously presented. Time point/electrode combinations no included in a significant cluster are shown in gray; for those included in a significant cluster, the color indicates the *F* statistic at each location. Note that the cluster shown for the Neutral vs New effect is only marginally significant (*p* =.059).

For trials given a Guess or New judgment, the valence effect was not significant (*p*s > .40; see waveforms and scalp maps are available in the Supplementary Materials at https://osf.io/jyd5c/).

To characterize the general time course of memory retrieval (i.e., old/new effects) independent of emotion effects, we compared words previously paired with a neutral picture (face or scene) and given a Remember or Know judgment to words that had not been presented at encoding and given a Guess or New judgment. This analysis allowed comparisons between the timing of emotion effects and general memory effects to directly assess our second research question. As can be seen in Figure 1, previously presented words elicited a more positive amplitude from around 400 to 700 ms at frontal electrodes, and from around 500-750 ms at parietal electrodes. A cluster from 408-721 ms spanning a large range of electrodes was marginally significant (*p* = .051).

The full results of all mass univariate analyses are available in the supplementary materials at https://osf.io/jyd5c/.

## Discussion

In the current study we sought to investigate the electrophysiological correlates and time course of memory retrieval for neutral words previously encoded in a neutral, positive, or negative context. Using ERPs we found evidence that, compared to a neutral encoding context, words encoded in a positive or negative context began to show a more positive-going amplitude in parietal electrodes beginning around 200 ms post stimulus onset. This occurred before the general old-new recognition effect, which began around 400 ms.

### Early effects of emotional encoding context at retrieval

Theories of memory suggest that processes engaged at encoding are reactivated at retrieval, and that this recapitulation plays an important role in determining retrieval success (Morris et al., 1977; Tulving & Thomson, 1973). In line with such views, a large empirical literature using fMRI shows substantial overlap in neural activation during encoding and retrieval (for reviews see Bowen et al., 2018; Buckner & Wheeler, 2001). However, the temporal resolution of fMRI means that it is unclear whether these reactivations occur before, during or after successful retrieval. Although the EEG literature on this question is still small, a few studies have suggested that neural recapitulation can be observed quite early during retrieval (Johnson et al., 2008, 2015; Waldhauser et al., 2016).

In previous fMRI work, including a study using the same paradigm employed in the present study, our lab has shown that neural recapitulation is greater for negatively valenced stimuli than neutral or positive (Bowen & Kensinger, 2017a, 2017b). Only a few studies have examined how emotional content from encoding affects the ERP response at retrieval (Jaeger et al., 2009; Maratos & Rugg, 2001; Smith, Henson, Dolan, & Rugg, 2004). Although these studies have suggested that emotion effects may emerge early during retrieval, these effects have been inconsistent and most of these studies have not examined both negative and positive valence.

The present results showed that encoding a neutral word in an emotional context—either positive or negative—can affect the neural response from the very earliest stages of memory retrieval. Indeed, the effect of emotion in our data began around 200 ms after the onset of the neutral retrieval cue. Not only is this coincident with the earliest old/new differences generally observed in ERP (Wilding & Ranganath, 2011), it is earlier than we were able to distinguish words encoded in neutral contexts from new words in the present data. This suggests that the emotion of the encoding context is reactivated at the earliest stages of memory retrieval and may play a role guiding the retrieval process.

It is not entirely clear what neural process is represented by the emotion effect we observed. One interpretation is that it reflects modulation of the left parietal old/new effect that has been previously associated with memory recollection in a large ERP literature (Wilding & Ranganath, 2011). Some work has suggested that emotion of the encoding context can modulate this component (Maratos & Rugg, 2001), and the effect of emotion in the present study showed a scalp distribution that is consistent with this effect (see Figure 1). Under this interpretation, participants showed greater episodic recollection for items that had been encoded with an emotional image, perhaps because the emotional properties of the image led it to be encoded and then recalled in greater detail. Alternatively, the effect we observed could be interpreted as the late positive potential (LPP) that is generally observed to emotional stimuli, and which appears as a very similar parietal positivity (Hajcak, Weinberg, MacNamara, & Foti, 2011). This interpretation would be in line with a more direct recapitulation account: it would suggest that the same neural processes that are observed when emotional stimuli are being viewed can be elicited by the incidental retrieval of emotional stimuli, even when the retrieval cue is neutral.

Under either of these interpretations, the timing of the effect is somewhat surprising. The left parietal old/new effect generally begins around 500 ms and lasts a few hundred milliseconds. If the present effect is identified with this component, it would suggest that emotional contexts can lead retrieval processes to begin earlier than they are usually observed. The LPP generally begins between 300 and 500 ms in response to emotional pictures and usually lasts at least a few hundred milliseconds, but can last longer depending on the task (Gable, Adams, & Proudfit, 2015). If the present effect is identified with this component, it suggests that cued retrieval of emotional information can be faster than initial recognition of the emotional properties of stimuli.

Of note, even though we report emotion-related differences in the electrophysiological signature, we did not find any behavioral differences in memory performance as a function of encoding context. This is not surprising given the short delay between encoding and retrieval blocks. Most findings of an emotion effect on memory have retention intervals upwards of 24 hours to allow for consolidation processes to unfold (Sharot & Yonelinas, 2008). Further, the to-be-remembered stimuli were not emotional themselves, but were neutral words, and participants were never explicitly asked to associated the word and emotional context, nor make a judgment about the emotionality of the encoding stimuli at any point during the experiment. These procedural details likely contributed to no differences in behavior, but also made for a strong test of emotion effects on memory reactivation.

### Effects of negative vs. positive valence

Notably, when the present paradigm was employed in fMRI, there was greater recapitulation in sensory processing regions during recollection of words encoded in a negative context compared to positive or neutral (Bowen & Kensinger, 2017b). In the ERP results, there was no evidence of greater reactivation for negative context specifically; instead, negative and positive valence showed an equally increased positivity compared to neutral, consistent with the ERP findings of Smith et al. (2004). Although ERP and fMRI are complementary techniques, they are sensitive to different aspects of the neural response for a number of reasons (Ekstrom, 2010; Lau, Gramfort, Hämäläinen, & Kuperberg, 2013; Logothetis, 2008; Murta, Leite, Carmichael, Figueiredo, & Lemieux, 2015). In the present case, it is likely that the timing of the visual activity at retrieval was variable across trials. Because the time course of the BOLD signal in MRI is slow and because the processing of MRI data generally presents activation that is a weighted average across several seconds of data, this variability would make little difference to the MRI results. However, in ERP, the visual response consists of a series of fast deflections of opposite electrical polarity (Pratt, 2011). If these are not aligned precisely in time across trials, they are likely to cancel out when averaging across trials. Thus, the ERP response may have captured broader effects of emotion present in both positive and negative conditions, while missing sensory-specific valence differences. Future work using time-frequency analysis methods that ignore phase and/or methods that make sensory recapitulation more clearly identifiable may be able to use the temporal resolution of EEG to examine the time-course of valence modulated sensory recapitulation (see, for example, Waldhauser et al., 2016, who were able to show rapid recapitulation of visual activity by combining time-frequency analysis with lateralized presentation at encoding).

### Limitations and Future Directions

One interesting question is whether the emotion effects on reactivation are stronger for recollected compared to familiar stimuli. Prior ERP work has shown that reactivation effects were enhanced for remember compared to know responses (Johnson et al., 2008) and using fMRI we also found evidence that there was greater recapitulation for remember compared to know responses, suggesting the reactivation may be more strongly associated with recollection processes (Bowen & Kensinger, 2017b). We broadened our ERP analyses to trials given either a remember or know response, rather than comparing effects for these response types, because we did not have enough trials to adequately assess this. The ability to examine remember and know trials separately may also help to determine the relationship of the emotion effect we observed to the left parietal old/new effect, since this effect is generally associated specifically with recollection (i.e., remember trials). On the other hand, if the effect we observed is a recapitulation of the emotional LPP effect, this effect could appear in the absence of recollection if the neutral cues have been imbued with some of the emotional properties of the pictures they were paired with (cf. Jaeger et al., 2012).

A limitation of our encoding procedure was the placement of target words above the picture. As a result of this stimulus placement, the EEG and ERPs at encoding indicated that participants were switching their gaze from the word to the picture in a way that was not consistent across trials and subjects. This resulted in difficulty time-locking the ERP to a particular stimulus or process at encoding, thus encoding data were not analyzed. Future work with encoding procedures that are better able to streamline participant attention to both stimuli simultaneously, perhaps with one stimulus superimposed on the other (re: Jaeger & Rugg, 2012) or the stimuli following each other temporally, would allow for a more detailed analysis of recapitulation by allowing for direct comparison of similarity in the EEG at encoding and retrieval.

## Conclusions

The study of emotion on memory has primarily focused on encoding processes, with little emphasis on how emotion might influence downstream processes, including retrieval. Prior fMRI work has revealed that emotion modulates the patterns of reactivation during successful memory reactivation, but does not reveal the time course of these effects. The current ERP findings show that effects of emotion at retrieval are not simply a result of successful retrieval. Instead, our findings show that effects of the emotionality of the encoding context are present very early during the retrieval process and thus may play a role in guiding successful retrieval.

## Funding

This work was supported by a National Institute of Health Grant awarded to EAK [R01MH080833].

## Footnotes

1 The electrodes were: Fp1, AF3, F7, F3, FC1, FC5, T7, C3, CP1, CP5, P7, P3, Pz, PO3, O1, Oz, O2, PO4, P4, P8, CP6, CP2, C4, T8, FC6, FC2, F4, F8, AF4, Fp2, Fz, Cz

2 The guess response was instructed to be used when participants were unsure whether the stimulus was old or new. Given this ambiguity, these trials were not included in any behavioral analysis.

